# Identifying drivers of sewage-associated pollutants in pollinators across urban landscapes

**DOI:** 10.1101/2023.01.20.524979

**Authors:** Michael F. Meyer, Matthew R. Brousil, Benjamin W. Lee, Madison L. Armstrong, Elias H. Bloom, David W. Crowder

## Abstract

Human sewage can introduce pollutants into food webs and threaten ecosystem integrity. Among the many sewage-associated pollutants, pharmaceuticals and personal care products (PPCPs) are useful indicators of sewage in ecosystems and can also cause potent ecological consequences even at minute concentrations (e.g., ng/L). Despite increased study over the past three decades, PPCPs in terrestrial systems have been less studied than those in aquatic ecosystems. To evaluate PPCP prevalence and drivers in a terrestrial ecosystem, we analyzed managed and native bees collected from agroecosystems in Washington State (USA) for PPCPs. Caffeine, paraxanthine, cotinine, and acetaminophen were detected in all three evaluated taxa (*Bombus vosnesenskii, Agapostemon texanus*, and *Apis mellifera*), with *B. vosnesenskii* and *A. texanus* having a higher probability of PPCP detection relative to *A. mellifera*. The probability for PPCP presence in all three taxa increased in landscapes with more human development or greater plant abundance, with significant but negative interactions among these factors. These results suggest that human activity, availability of resources, and species-specific traits affect the introduction and mobilization of PPCPs in terrestrial ecosystems. Consequently, monitoring PPCPs and their ecological responses in terrestrial ecosystems creates opportunities to synthesize consequences of sewage pollution across terrestrial and aquatic ecosystems and organism types.

## INTRODUCTION

The introduction of wastewater and its byproducts into ecosystems can mobilize pollutants that reshape communities and food webs (Edmondson 1970). Historically, research on wastewater pollution has largely focused on changes in effluent nutrient concentrations as well as chemical and biological oxygen demand (Edmondson 1970; Brydon and Frodsham 2001; Tong et al. 2020). Recently, research emphasis has broadened to include micropollutants that are often found in sewage (Bernhardt et al. 2017). Pharmaceuticals and personal care products (PPCPs) in particular have garnered increased attention as an emerging organic micropollutant because they are consistently associated with human sewage and pose potent, yet often uncertain, ecological consequences (Richmond et al. 2017; Meyer et al. 2019).

As PPCPs are consistently associated with sewage, their presence, even at minute concentrations (e.g., ng/L), can indicate wastewater inputs into an ecosystem. Previous continental and worldwide surveys show that PPCPs such as antibiotics, non-prescription and prescription drugs, hormones, and fragrances are pervasive within subsurface and surface systems (Kolpin et al. 2002; Focazio et al. 2008; Wilkinson et al. 2022). Once introduced into ecosystems, PPCPs can propagate through food webs, where they can be metabolized (del Rey et al. 2011), accumulate within organisms (Meador et al. 2016), and be transferred across trophic levels (Richmond et al. 2018). However, patterns of PPCP prevalence across taxa can be related to individual behavior and tissue allocation (Meador et al. 2016), and biological responses to PPCPs are often uncertain and context-dependent. For example, activity and feeding rates of individual fish (*Perca fluviatilis*; European perch) increased when they were exposed to the anti-anxiolytic drug oxazepam, but sociality of fish populations decreased (Brodin et al. 2013). Certain algal taxa have been shown to have reduced photosynthesis and increased 16-carbon unsaturated fatty acid production when exposed to fluoxetine, even though essential fatty acid synthesis was not affected (Feijão et al. 2020). Bumble bees fed nectar primed with caffeine have shown increased foraging behavior on nectars with similar aromatic compounds (Wright et al. 2013; Arnold et al. 2021). Overall, the literature suggests that biological responses to PPCPs are common and often deleterious, although specific responses may also be uncertain (Richmond et al. 2017).

Although considerable evidence from aquatic systems suggests that PPCPs are ubiquitous and disrupt ecological processes, comparatively few studies on PPCPs have been conducted in terrestrial systems (Meyer et al. 2019). This imbalance in the literature creates opportunity to assess how PPCPs may propagate through terrestrial food webs in comparison to aquatic food webs. For example, bees are exclusively terrestrial insect taxa that provide pollination services in natural and managed ecosystems (Kleijn et al. 2015). As has been observed in earthworms (Carter et al. 2021), bee pollinators may be commonly exposed to PPCPs through soil contact, interactions with plants, applications of biosolids for fertilizer, or through contamination of water in terrestrial ecosystems. Due to the dramatic, recent declines in bee pollinators within terrestrial food webs worldwide (Potts et al. 2010; Kleijn et al. 2015), bee taxa may be promising, societally important model organisms for expanding PPCP research from aquatic to terrestrial ecosystems.

To assess PPCP prevalence and mobilization in pollinators within exclusively terrestrial food webs, we evaluated the presence and drivers of PPCPs in three bee species: *Bombus vosnesenskii, Agapostemon texanus*, and *Apis mellifera*. Two of these species are wild (*B. vosnesenskii, A. texanus*), but *A. mellifera* (honey bee) is managed by humans. Our first goal was to identify species-specific patterns of PPCP presence. We predicted that taxa interacting with the soil (ground-nesting bees; i.e., *B. vosnesenskii, A. texanus*) would be more frequently associated with PPCPs than managed, colony-forming species (i.e., *A. mellifera*), as ground-nesting taxa would more likely encounter PPCPs within groundwater and biosolids, similar to interactions observed in earthworms (Carter et al. 2021). Our second goal was to assess the potential drivers of PPCP presence in each bee taxa. We predicted that PPCP presence would increase at sites with greater human development and with a higher density of floral plant resources at the sampling location, both of which can be potentially concentrated sources of PPCPs (Carter et al. 2021; Meyer et al. 2022). Overall, our study provides some of the first evidence for PPCP uptake in exclusively terrestrial bee species, while also demonstrating that both landscape-context and organismal traits may affect exposure to and uptake of PPCPs.

## MATERIALS AND METHODS

### Study System

Our study of PPCPs in bees was nested within a study of bee ecology on 36 small (< 25 ha), diversified farms and community gardens (e.g., farms and gardens with more than five unique flowering crops in simultaneous production) in western Washington (USA) (Bloom et al. 2021). For the present study, bees were collected from a subset of 10 urban gardens that were selected along an urbanization gradient (Fig. 1). Each garden was at least 1 km apart for spatial independence (Bloom et al. 2021; Desaegher et al. 2022).

**Figure 1:**
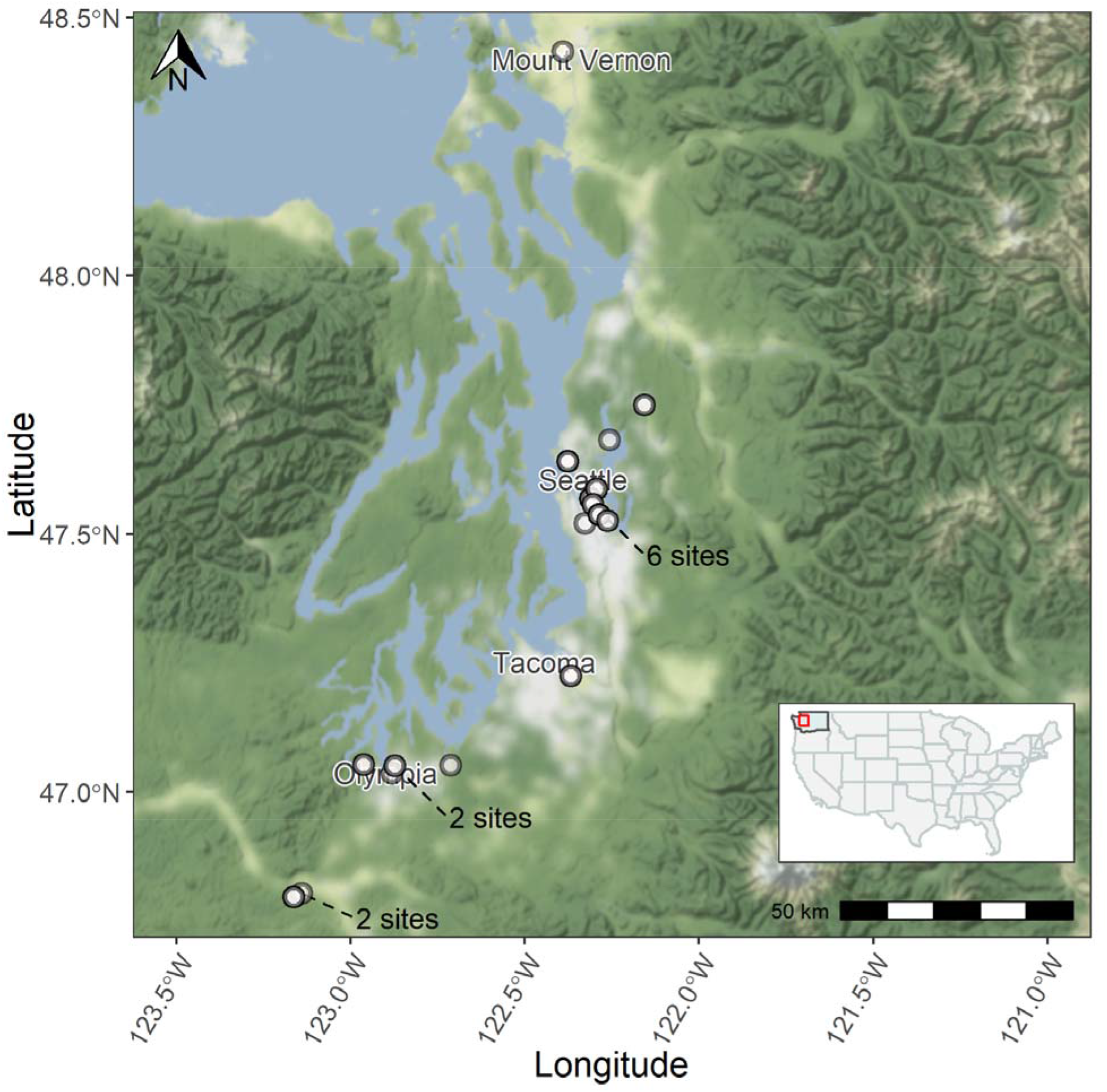
Map of bee sampling locations (white points). Map tiles by Stamen Design (stamen.com), under CC BY 3.0 (creativecommons.org/licenses/by/3.0). Data by OpenStreetMap (openstreetmap.org), under ODbL (openstreetmap.org/copyright). State data from US Census Bureau 2019.

### Bee and plant surveys

Bees were collected at each site using three blue vane traps (SpringStar LLC, Woodinville, WA, USA) and 15 bee bowls placed along a 50 m transect. All collected bees were preserved, pinned, and then identified to species (Bloom et al. 2021). Of the 6,539 specimens collected, we selected 101 specimens with approximately balanced specimens from each of the three bee species (*B. vosnesenskii*, *A. texanus*, and *A. mellifera*); these taxa reflect variation in social behavior (social vs. solitary taxa) and nesting strategies (i.e., cavity vs. ground-nesting taxa). Because bee species were collected across three years (2014, 2015, 2016) and three time points (spring, summer, and fall), we selected pinned specimens for this analysis to have a balanced number of individuals across the sampling period. Specimen selection was stratified by species, year, early/mid/late-summer timing, and sampling location, where each species-timing-year-site combination included at least two individuals. As each species-timing-year-site combination contained numerous individuals from the original sampling events, personnel haphazardly selected individuals within a species-timing-site-year stratum.

To assess the role of plant communities in affecting PPCP presence in bees, plant richness and abundance were measured at sites on the same date as bee sampling. Plants with flowers serving as resources for bee visitation (i.e., having pollen and nectar) were recorded along the same transect used for bee collection. A portable 1 × 1 m plot was placed over vegetation at 5 m intervals and all plants with open flowers were identified to species. Each transect moved in a serpentine fashion across each study site. Summarized field variables are detailed in Table S1.

### Landscape context

As sites were located along an urbanization gradient, we determined the amount of developed landscape within 1 km of each site using the United States Department of Agriculture cropland data layer (CDL). Proportion development was defined as the count of developed pixels within 1 km of each site divided by the total number of all pixels. Each pixel within the CDL classifies a 30 × 30 m area as a landscape class, with 255 possible classifications. To characterize all human development within a landscape, we summed across low, moderate, and high intensity pixels (Table S1) (Han et al. 2014).

### PPCP extraction and quantification

Each dried bee sample was massed and then ground with mortar/pestle. Bee PPCPs were extracted using a three-phase sequential extraction, similar to those described previously in Brodin et al. (2013) and Furlong et al. (2008). First, 1.5 mL of methanol/water mix (7:3 ratio) with 0.1% formic acid was added into the mortar with the bee parts and poured into a glass culture tube. An additional 1.5 mL methanol/water mix with 0.1% formic acid was used to rinse the mortar and pestle, and then was poured into the same centrifuge tube. Tubes with tissue and extract were centrifuged at 2,000 rpm for 5 minutes. Following centrifugation, supernatant was then added to a clean centrifuge tube, and then immediately wrapped in parafilm. A second 1.5 mL acetonitrile was next added to the first tube containing the bee tissue. The tube containing bee tissue was then vortexed and centrifuged twice. After each of the three extraction phases, the sample was placed under nitrogen flow in a 40°C bath. Once samples were nearly evaporated completely, 1mL of formate buffer was added to each sample, and the concentrated extract was transferred to a 1.5 mL amber glass autosampler vial. Samples were preserved in the dark at −20°C until being analyzed with HPLC/MS.

### HPLC/MS Quantification

PPCP identification and enumeration followed methods similar to Furlong et al (2008) and Brodin et al. (2013). Standards for each PPCP are described in table SM1. HPLC eluents included a 10-mM formate buffer and 100% acetonitrile solution that varied in percent contribution over the quantification procedure (Table SM1). Minimal detection limits were estimated to be 5 ng/g. A main difference between our methods and others prior is that we split analyte signatures into separate channels, so as to avoid peak interference (described in Table SM1). Mass spectrometer time-programmed operating conditions for individual compounds are detailed in Table SM2-3.

Samples were also analyzed in a manner to account for potential cross-sample contamination and peak drift. Following standard samples, 2 blanks of 100% methanol were processed and assessed for contamination. Following the two blank samples, bee samples were processed in batches of 10 followed by 1 blank, 1 standard (100 ng/L), and 1 blank. This routine allowed us to purge the column from potential cross-sample contamination, to assess if samples were contaminating downstream samples within a batch, and to control for peak drift.

Following peak quantification, we noticed that concentrations tended to be bimodal, where PPCPs were either detected in higher concentrations or not detected (Figure 2). We assumed that the bimodality of these detections was likely a product of low-to-intermediate PPCP concentrations degrading since the time of specimen collection or diffusing from bee tissues as samples were originally collected in an ethanol solution before pinning (Bloom et al., 2021). In order to provide conservative estimates of bee PPCP presence, we reduced concentrations into categorical presence/absence informatics, such that subsequent models and model interpretations should be conservative.

**Figure 2:**
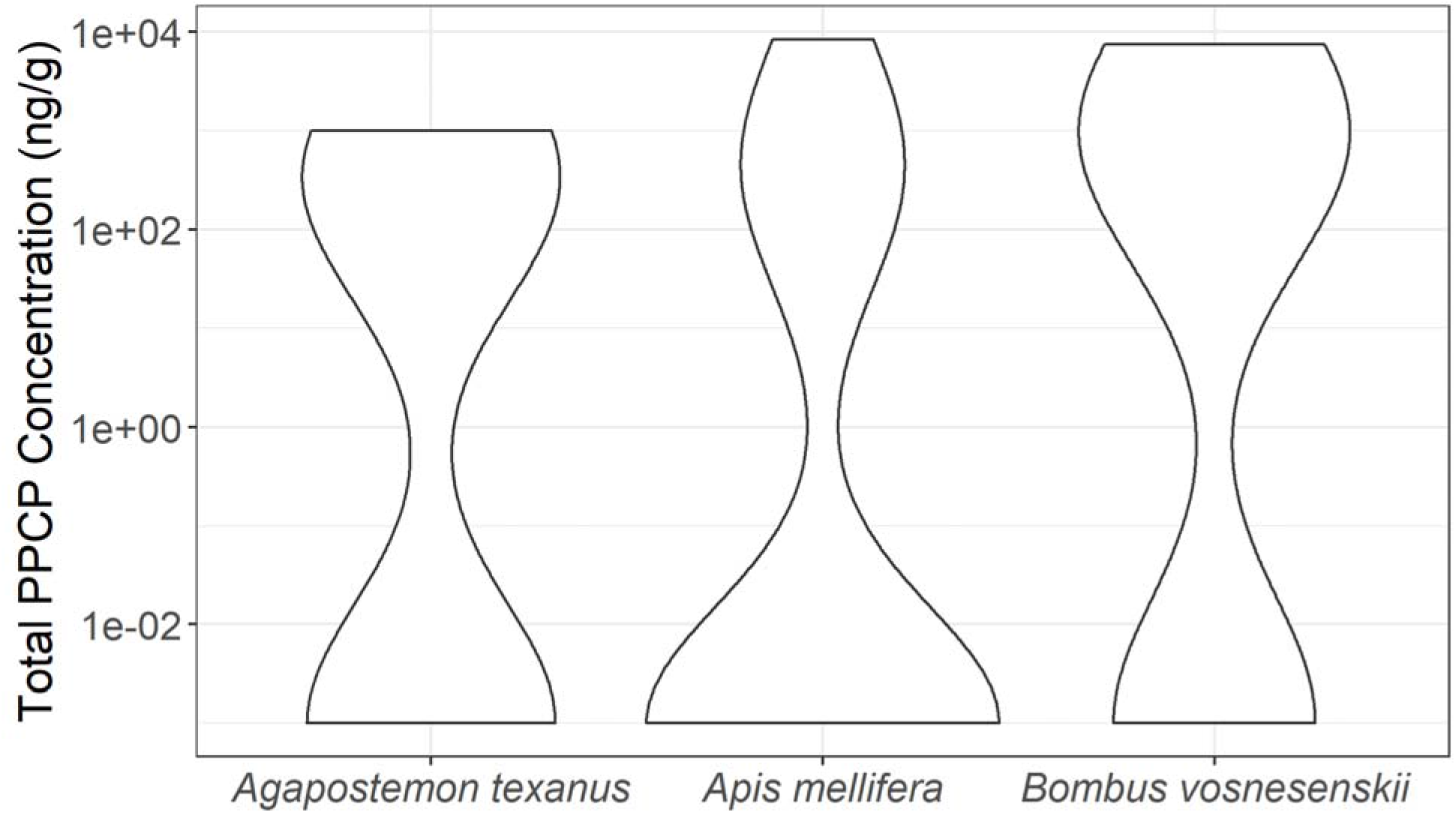
Violin plot of total PPCP concentrations (on log10 scale) in each of the three examined bee taxa. Although the figure shows continuous PPCP concentrations, few samples contained “intermediate” PPCP concentrations, meaning that concentrations were either high (i.e., greater than 500 ng/g) or not detectable. Given that this framework seemed unrealistic and could be a product of how bees were preserved, our main analysis only focused on presence/absence of PPCPs in samples. Therefore, our logistic regression approach should be more conservative than continuous, linear analyses. Nevertheless, trends in PPCP concentrations mirror patterns observed in PPCP presence/absence, where *B. vosnesenkii* and *A. texanus* have higher concentrations and probabilities of PPCP presence relative to *A. mellifera*.

**Figure 2:**
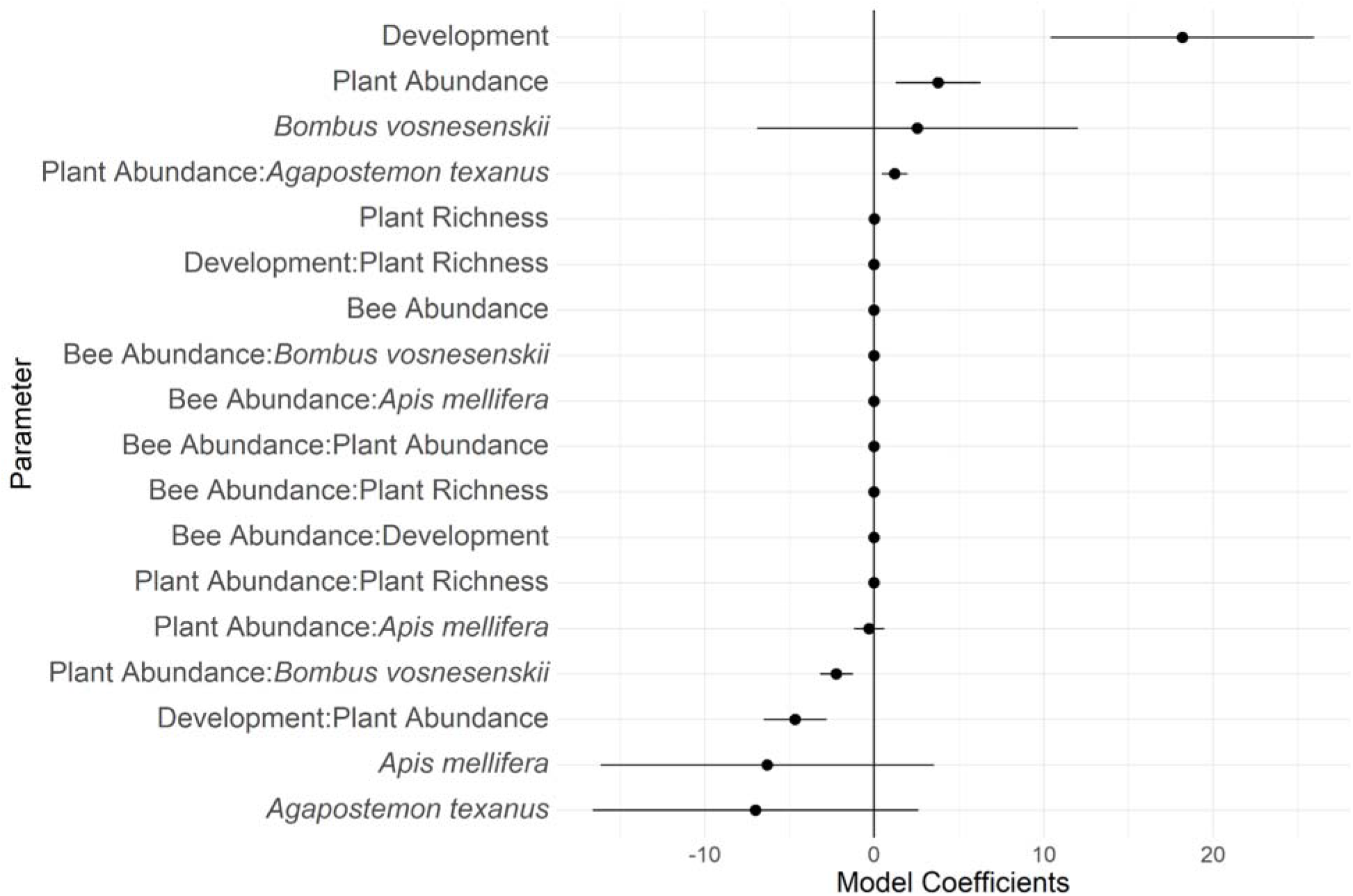
Coefficients for the final averaged model using all data for training. Points indicate th**e** estimated model coefficient, and bars reflect the adjusted standard error. A value of 0 indicates **a** variable does not discriminate between PPCP presence or absence. Values greater or less than zero correspond to predictors that are more or less likely to be associated with PPCP presence, respectively. Parameters with a colon indicate two-way interaction terms. Parameters are arranged by decreasing coefficient estimate value.

### Quality Assurance and Quality Control Procedures

During PPCP extraction and quantification, an internal standard was unintentionally not included within each sample. To account for potential biases and contamination that might arise during the PPCP extraction and quantification process, we developed quality assurance and quality control protocols to ensure high confidence of model results.

Throughout the entirety of the sampling process, laboratory personnel refrained from consuming caffeinated beverages, nicotine products, and non-prescription medications. Personnel wore an N95 mask and nitrile gloves during extraction to reduce chance of contamination during extraction.

Samples were double-blind throughout the entire specimen selection, PPCP extraction, peak quantification, and modeling procedures. Personnel extracting PPCPs from bees were only aware of “Sample ID” for each specimen, which was written on the autosampler vial. Experimental autosampler vials were haphazardly placed in the auto-sampler to reduce likelihood of biases due to order in the auto-sampler. A separate laboratory personnel analyzed ion spectra. Throughout the entire spectra analysis, the technician only knew the “Sample ID” and not any metadata associated with that “Sample ID”. Lastly, a third person modeled the data from HPLC/MS quantification. Through each step, experimental personnel only knew samples by a unique “Sample ID” and were not aware of any attributes associated with that sample. Together, this scheme was intentionally designed to potentially eliminate personnel-specific biases throughout the extraction and analysis process.

### Modeling PPCP presence

We iterated through all combinations of logistic regression models to predict PPCP presence based on six predictor variables: bee and plant abundance, bee and plant diversity, percent development, bee taxon, and all two-way interactions between these variables (Katz et al. 2015; Rheubert et al. 2020). To account for edge effects, all variables were standardized by the perimeter:area ratio of their respective site. Once all models were generated, we selected the best model based on AICc, R^2^, AUC, accuracy (percent true results), and p-value. When multiple models had similar performance (AICc within two points of lowest AICc value and R^2^, AUC, and accuracy above the median of all models), best performing models were averaged to create a single statistical model. To validate our results, we repeated analyses 1,000 times with 80:20 train:test subsetted data, and compared the distribution of model parameters and pseudo-R^2^ values from subsetted data to those models generated with the entire dataset. This subsetting routine allowed us to assess whether the distribution of possible model parameters was multi-modal and to evaluate whether the model constructed with 100% of the training data was overfit to those data (Rheubert et al. 2020).

All analyses were conducted within the R Statistical Environment (R Core Team 2022) using the packages glmulti (Calcagno 2019), tidyverse (Wickham et al. 2019), janitor (Firke 2020), MuMIn (Barton 2020), ggpubr (Kassambara 2019), ggeffects (Lüdecke 2018), lubridate (Grolemund and Wickham 2011), plotrix (Lemon 2006), Hmisc (Jr et al. 2020), corrplot (Wei and Simko 2017), ggrepel (Slowikowski 2019), ggspatial (Dunnington 2021), tigris (Walker 2021), cowplot (Wilke 2019), sf (Pebesma 2018), and readxl (Wickham and Bryan 2019).

## RESULTS

### Species-specific PPCP detections

We detected four PPCPs across all three bee species: caffeine, paraxanthine/1,7-dimethylxanthine, acetaminophen/paracetamol, and cotinine. We did not detect evidence of seven other PPCPs: codeine, warfarin, trimethoprim, sulfamethoxazole, diphenhydramine, thiabendazole, or albuterol. *Bombus vosnesenskii* (50% of samples; Table S2) and *A. texanus* (44%; Table S2) tended to have a higher odds ratio of PPCP presence relative to *A. mellifera* (26%; Table S2).

### Relating PPCP presence with human development and plant abundance

Across all bee taxa, the probability of PPCP presence increased in landscapes with greater human development or higher plant abundance (Fig. 2). However, there was a significant negative interaction between human development and plant abundance, suggesting the positive association between development and PPCP presence decreased in landscapes with greater plant abundance; and, the positive effects of plant abundance on PPCP presence decreased in landscapes with greater development (Fig. 2, Fig. S1).

Our subsetting routine, which assessed the probability of observing the final model solely by chance, suggested our final model coefficients were non-random (Fig. 3). Model parameter estimates (Figure 3) as well as pseudo-R^2^ values (Figure 4) were generally unimodal. Most subsetted model runs included human development and plant abundance as positive predictors of PPCP presence (Figure 3), and most coefficients for interactions between plant abundance and development were negative (Figure 3). Coefficients from the subsetting routine also corresponded with patterns of PPCP detections for each taxon, where *B. vosnesenskii* had higher positive coefficients, and *A. mellifera* and *A. texanus* tended to have lower negative coefficients (Figure 3).

**Figure 3:**
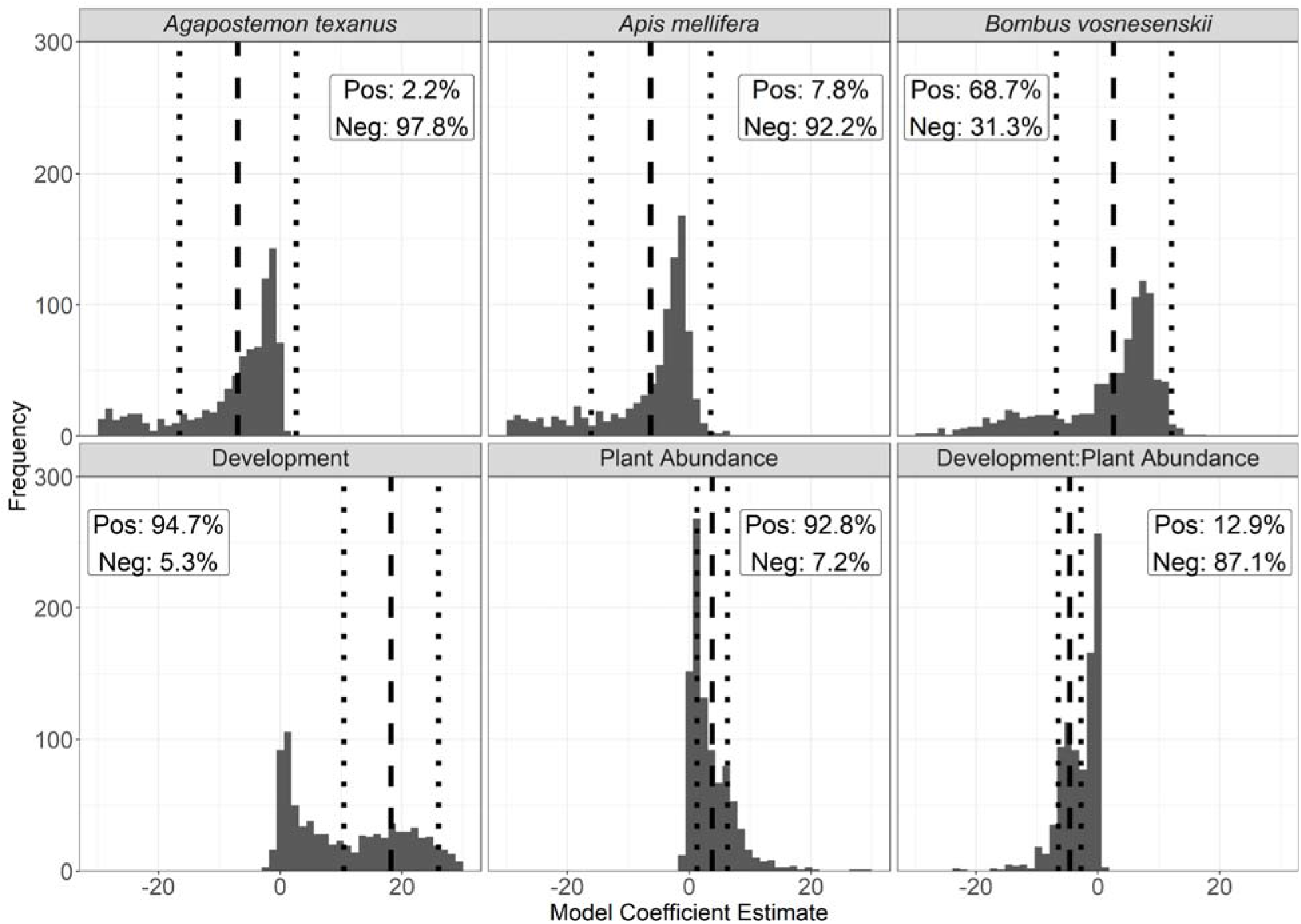
Distributions of most influential model coefficients for the final averaged model using 80% of original data as training data. Model averaging protocols were repeated 1,000 times, with each iteration starting with a species-stratified, random subsample. Vertical, dashed lines indicate the coefficient of the parameter that appeared in the final averaged model, when 100% of the data were used. Vertical, dotted lines indicate the adjusted standard error of the coefficient of the parameter that appeared in the final averaged model, when 100% of the data were used. Labels within each facet reflect the percent of coefficients that were positive (i.e., greater than zero) and negative (i.e., less than zero).

**Figure 4:**
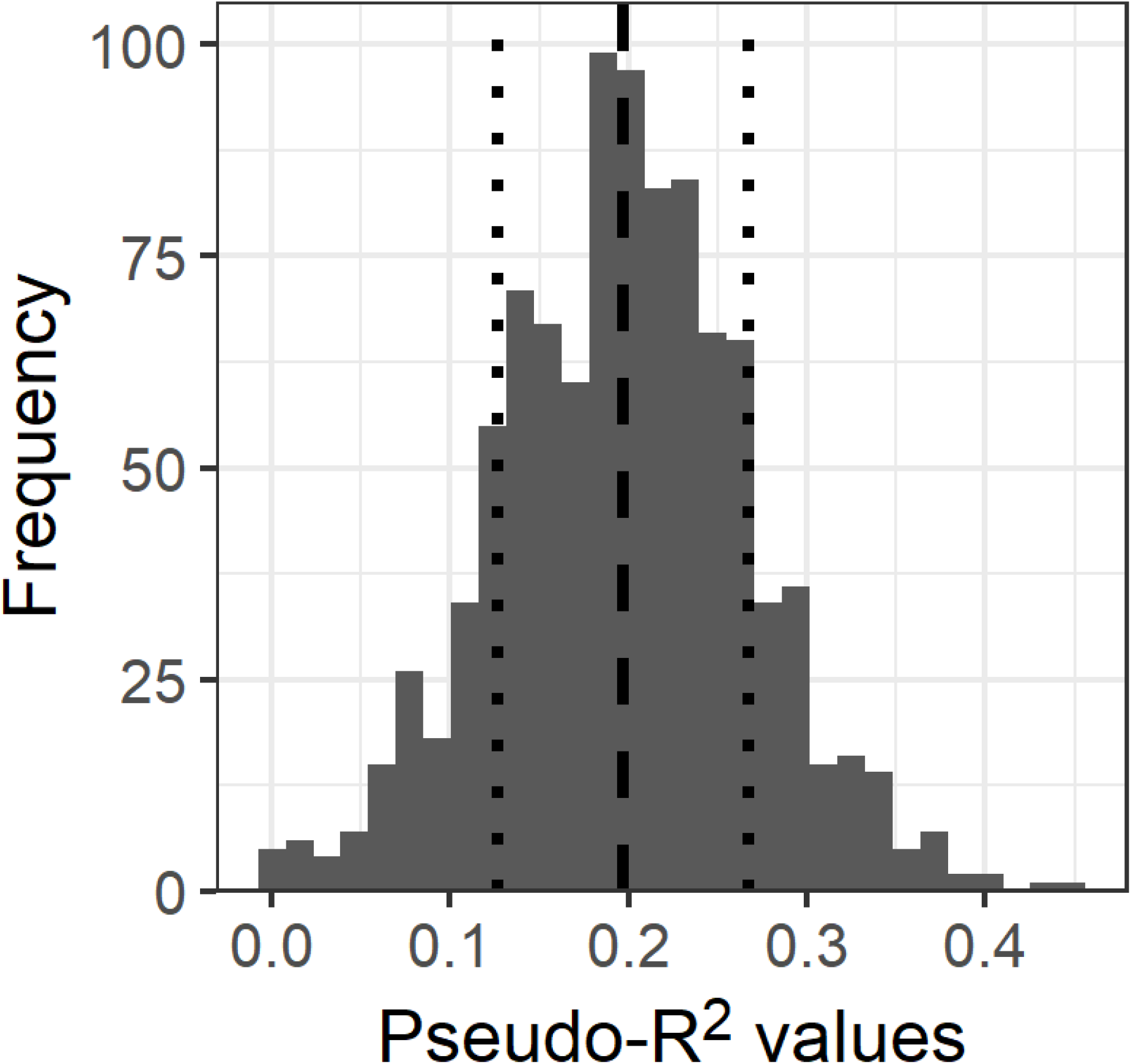
Distribution of pseudo R^2^ values from all averaged models in the permuted analysis. The vertical dashed line represents the mean of the distribution (Pseudo R^2^ = 0.197), and vertical, dotted lines represent one standard deviation from the mean (sd = 0.07).

## DISCUSSION

Our study demonstrates the presence of PPCPs in a terrestrial ecosystem and highlights that pinned insect specimens can be used along with environmental data to understand patterns of PPCP prevalence. These results are notable, considering that the study of PPCPs in terrestrial ecosystems is uncommon relative to aquatic environments (Meyer et al. 2019). Our results corroborate those results from aquatic systems, showing that PPCP presence can be influenced by human development and species-specific traits (Bendz et al. 2005; Meador et al. 2016) and that PPCPs can be mobilized in terrestrial food webs (Lagesson et al. 2016; Richmond et al. 2018).

Our results suggest PPCPs are more prevalent in *B. vosnesenskii* and *A. texanus* than in *A. mellifera* (Table S2), implying that differences in species’ life history can affect PPCP accumulation. Both *B. vosnenskii* and *A. texanus* create nests in the soil, which may increase exposure to human contaminants in the soil matrix and groundwater relative to *A. mellifera*, which is managed in artificial nests by humans (Gradish et al. 2019). Additionally, *A. mellifera* may prefer different resources within the same habitat (Thompson and Hunt 1999; Leonhardt and Blüthgen 2012), generating variation in exposure. Niche partitioning of floral resources due to differences in traits, such as tongue length, could also mediate pollutant exposure (e.g., pesticides) (Brittain and Potts 2011). In addition to pollinator traits, heightened PPCP exposure for *B. vosnenskii* and *A. texanus* may be due to variable production practices across sampling locations, with famer’s application of wastewater byproducts to soil contaminating certain pollinator nesting sites (Karnjanapiboonwong et al. 2010; Shahriar et al. 2021). While we are not aware of wastewater byproduct applications at our sampling locations, prevalence of external pollutants in conventional and organic farms is common (Humann-Guilleminot et al. 2019).

Beyond species-specific patterns, our results suggest that human development and plant abundance mediated PPCP presence across all three taxa. This result is consistent with previous findings in aquatic systems, where the source concentration (e.g., amount of development) and paths to trophic transfer (e.g., number of plants) are correlated with the presence of PPCPs at higher trophic levels (Richmond et al. 2018). For example, PPCP concentrations in aquatic systems have been shown to be directly proportional with human population size and inversely proportional to distance from a human population center (Bendz et al. 2005; Meyer et al. 2022). PPCPs also can transfer between trophic levels (Richmond et al. 2018), which may lead to non-linear biological processes and consequences, such as our data showing a negative interaction between plant abundance and human development. Consequences of PPCPs within food webs are hard to predict, although they can cause biological responses at physiological (Feijão et al. 2020), behavioral (Brodin et al. 2013), population (Hoppe et al. 2012), community (Lee et al. 2016), and ecosystem (Richmond et al. 2019) levels. Aside from PPCPs eliciting direct biological responses, PPCPs may co-occur with numerous human disturbances, such as nutrient pollution, that may further obfuscate clear associations between biological consequences and PPCP exposure.

To our knowledge, this is the first study to detect PPCPs in bee tissues that were preserved and pinned in a manner similar to museums. While not originally intended for the present study, detailed field notes and extensive covariate collection created the opportunity for us to evaluate PPCP presence across taxa while testing potential mechanistic drivers of PPCP presence. Considering the growing number of pharmaceuticals (Daughton and Ternes 1999) and changes in pesticides used on national and global markets (Douglas et al. 2020), preserved collections, such as in natural history museums, may offer ripe and previously untapped opportunities to explore contaminant accumulation and mixtures within biota. For example, lake sediment cores have been used to reconstruct interdecadal PPCP contamination loadings (Anger et al. 2013). Similarly, museum collections may empower reconstructions of exposure, accumulation, and co-contaminant histories, all of which may be useful for detailing how contaminant loading and mixtures may change through time.

## Synthesizing PPCP patterns across ecosystem types

Over the past three decades, the study of PPCPs has expanded rapidly, and growing evidence suggests that PPCPs are pervasive micropollutants across aquatic and terrestrial systems alike. Our results suggest that much like aquatic systems, PPCPs tend to concentrate closer to their sources and have potential to enter food webs when vectors for trophic transfer are present. Additionally, our results demonstrate species-specific differences for PPCP uptake. Similar to patterns observed in PPCP accumulation in aquatic and riparian systems (Meador et al. 2016; Richmond et al. 2018), *B. vosnesenskii* and *A. texanus* had higher probabilities of PPCP detections relative to *A. mellifera*, implying that life histories, behavior, or physiological differences between taxa may play a role in PPCP uptake.

Broadly, our results present novel opportunities for assessing PPCP presence throughout food webs and suggest similarities to how terrestrial and aquatic systems accumulate PPCPs. Where PPCP data are rarer, pairing human development, environmental, and ecological data may aid managers in flagging systems that are more associated with PPCPs, and thus susceptible to declines in ecosystem function and services (e.g., pollination). Regardless of the exact trajectory, our study lays a foundation for future basic and applied PPCP research and creates a synoptic view of how organic contaminants may mobilize within aquatic and terrestrial environments.

## Supporting information

Supplemental tables and figures for main manuscript

Supplemental methods tables for main manuscript

## ACKNOWLEDGMENTS

We would like to thank S. E. Hampton, E. J. Rosi, and R. Spirits for offering advice on clarity and presentation in a previous version of this manuscript.

## AUTHOR CONTRIBUTIONS

MFM, MRB, BWL, EHB, and DWC designed the experiment. EHB collected field samples. MFM, BWL, and MRB developed scripts analyses and data visualization. MFM and MLA extracted PPCPs and analyzed HPLC/MS outputs. All authors contributed text, edited, and approved the final manuscript.

## FUNDING

Funding was provided to EHB and DWC by NSF GROW (121477-007), USDA ORG (2014-51106-22096), USDA Predoctoral Fellowship (2017-67011-26025), NSF Graduate Research Fellow (DEG 124006-001), Western SARE (GW15-022), and a grant from James Cook University, Cairns, QLD, AU.

## DATA AVAILABILITY STATEMENT

All data and scripts used to produce these analyses can be found in this project’s companion Open Science Framework repository (Brousil et al. 2022).

## Notes

https://osf.io/xyz8v/

### Competing Interest Statement

The authors have declared no competing interest.

## References

Anger, C. T., C. Sueper, D. J. Blumentritt, K. McNeill, D. R. Engstrom, and W. A. Arnold. 2013. Quantification of triclosan, chlorinated triclosan derivatives, and their dioxin photoproducts in lacustrine sediment cores. Environ. Sci. Technol. 47: 1833–1843. doi:10.1021/es3045289

Arnold, S. E. J., J.-H. Dudenhöffer, M. T. Fountain, K. L. James, D. R. Hall, D. I. Farman, F. L. Wäckers, and P. C. Stevenson. 2021. Bumble bees show an induced preference for flowers when primed with caffeinated nectar and a target floral odor. Curr. Biol. 31: 4127–4131.e4. doi:10.1016/j.cub.2021.06.068

Barton, K. 2020. MuMIn: Multi-Model Inference.

Bendz, D., N. A. Paxéus, T. R. Ginn, and F. J. Loge. 2005. Occurrence and fate of pharmaceutically active compounds in the environment, a case study: Höje River in Sweden. J. Hazard. Mater. 122: 195–204. doi:10.1016/j.jhazmat.2005.03.012

Bernhardt, E. S., E. J. Rosi, and M. O. Gessner. 2017. Synthetic chemicals as agents of global change. Front. Ecol. Environ. 15: 84–90. doi:10.1002/fee.1450

Bloom, E. H., E. C. Oeller, R. L. Olsson, M. R. Brousil, R. N. Schaeffer, S. Basu, Z. Fu, and D. W. Crowder. 2021. Documenting pollinators, floral hosts, and plant–pollinator interactions in U.S. Pacific Northwest agroecosystems. Ecology n/a: e3606. doi:10.1002/ecy.3606

Brittain, C., and S. G. Potts. 2011. The potential impacts of insecticides on the life-history traits of bees and the consequences for pollination. Basic Appl. Ecol. 12: 321–331. doi:10.1016/j.baae.2010.12.004

Brodin, T., J. Fick, M. Jonsson, and J. Klaminder. 2013. Dilute concentrations of a psychiatric drug alter behavior of fish from natural populations. Science 339: 814–815. doi:10.1126/science.1226850

Brousil, M. R., M. Meyer, B. W. Lee, M. L. Armstrong, E. H. Bloom, and D. Crowder. 2022. PPCPs in urban pollinators. Open Sci. Framew. doi:10.17605/OSF.IO/XYZ8V

Brydon, D. A., and D. A. Frodsham. 2001. A model-based approach to predicting BOD5 in settled sewage. Water Sci. Technol. 44: 9–16. doi:10.2166/wst.2001.0747

Calcagno, V. 2019. glmulti: Model Selection and Multimodel Inference Made Easy.

Carter, L. J., M. Williams, and J. B. Sallach. 2021. Uptake and Effects of Pharmaceuticals in the Soil-Plant-Earthworm System, p. 175–220. In S. Pérez Solsona, N. Montemurro, S. Chiron, and D. Barceló [eds.], Interaction and Fate of Pharmaceuticals in Soil-Crop Systems: The Impact of Reclaimed Wastewater. Springer International Publishing.

Daughton, C. G., and T. A. Ternes. 1999. Pharmaceuticals and personal care products in the environment: agents of subtle change? Environ. Health Perspect. 107: 907.

Desaegher, J., A. Ouin, and D. Sheeren. 2022. How far is enough? Prediction of the scale of effect for wild bees. Ecography n/a: e05758. doi:10.1111/ecog.05758

Douglas, M. R., D. B. Sponsler, E. V. Lonsdorf, and C. M. Grozinger. 2020. County-level analysis reveals a rapidly shifting landscape of insecticide hazard to honey bees *(Apis mellifera)* on US farmland. Sci. Rep. 10: 797. doi:10.1038/s41598-019-57225-w

Dunnington, D. 2021. ggspatial: Spatial Data Framework for ggplot2.

Edmondson, W. T. 1970. Phosphorus, nitrogen, and algae in Lake Washington after diversion of sewage. Science 169: 690–691.

Feijão, E., R. Cruz de Carvalho, I. A. Duarte, and others. 2020. fluoxetine arrests growth of the model diatom phaeodactylum tricornutum by increasing oxidative stress and altering energetic and lipid metabolism. Front. Microbiol. 11. doi:10.3389/fmicb.2020.01803

Firke, S. 2020. janitor: Simple Tools for Examining and Cleaning Dirty Data.

Focazio, M. J., D. W. Kolpin, K. K. Barnes, E. T. Furlong, M. T. Meyer, S. D. Zaugg, L. B. Barber, and M. E. Thurman. 2008. A national reconnaissance for pharmaceuticals and other organic wastewater contaminants in the United States - II) Untreated drinking water sources. Sci. TOTAL Environ. 402: 201–216. doi:10.1016/j.scitotenv.2008.02.021

Furlong, E. T., S. L. Werner, B. D. Anderson, and J. D. Cahill. 2008. Determination of human-health pharmaceuticals in filtered water by chemically modified styrene-divinylbenze resin-based solid-phase extraction and high-performance liquid chromatograph/mass spectrometry. Techniques and Methods 5-B5. Techniques and Methods 5-B5 US Geological Survey.

Gradish, A. E., J. van der Steen, C. D. Scott-Dupree, and others. 2019. Comparison of pesticide exposure in honey Bees (Hymenoptera: Apidae) and Bumble Bees (Hymenoptera: Apidae): Implications for risk assessments. Environ. Entomol. 48: 12–21. doi:10.1093/ee/nvy168

Grolemund, G., and H. Wickham. 2011. Dates and Times Made Easy with lubridate. J. Stat. Softw. 40: 1–25.

Han, W., Z. Yang, L. Di, A. L. Yagci, and S. Han. 2014. Making cropland data layer data accessible and actionable in GIS education. J. Geogr.

Hoppe, P. D., E. J. Rosi-Marshall, and H. A. Bechtold. 2012. The antihistamine cimetidine alters invertebrate growth and population dynamics in artificial streams. Freshw. Sci. 31: 379–388. doi:10.1899/11-089

Humann-Guilleminot, S., Ł. J. Binkowski, L. Jenni, G. Hilke, G. Glauser, and F. Helfenstein. 2019. A nation-wide survey of neonicotinoid insecticides in agricultural land with implications for agri-environment schemes. J. Appl. Ecol. 56: 1502–1514. doi:10.1111/1365-2664.13392

Jr, F. E. H., with contributions from C. Dupont, and many others. 2020. Hmisc: Harrell Miscellaneous.

Karnjanapiboonwong, A., A. N. Morse, J. D. Maul, and T. A. Anderson. 2010. Sorption of estrogens, triclosan, and caffeine in a sandy loam and a silt loam soil. J. Soils Sediments 10: 1300–1307. doi:10.1007/s11368-010-0223-5

Kassambara, A. 2019. ggpubr: “ggplot2” Based Publication Ready Plots.

Katz, S. L., L. R. Izmest’eva, S. E. Hampton, T. Ozersky, K. Shchapov, M. V. Moore, S. V. Shimaraeva, and E. A. Silow. 2015. The “Melosira years” of Lake Baikal: Winter environmental conditions at ice onset predict under-ice algal blooms in spring: Resolving Melosira years on Lake Baikal. Limnol. Oceanogr. 60: 1950–1964. doi:10.1002/lno.10143

Kleijn, D., R. Winfree, I. Bartomeus, and others. 2015. Delivery of crop pollination services is an insufficient argument for wild pollinator conservation. Nat. Commun. 6: 7414. doi:10.1038/ncomms8414

Kolpin, D. W., E. T. Furlong, M. T. Meyer, E. M. Thurman, S. D. Zaugg, L. B. Barber, and H. T. Buxton. 2002. Pharmaceuticals, hormones, and other organic wastewater contaminants in U.S. Streams, 1999-2000: A national reconnaissance. Environ. Sci. Technol. 36: 1202–1211. doi:10.1021/es011055j

Lagesson, A., J. Fahlman, T. Brodin, J. Fick, M. Jonsson, P. Byström, and J. Klaminder. 2016. Bioaccumulation of five pharmaceuticals at multiple trophic levels in an aquatic food web - Insights from a field experiment. Sci. Total Environ. 568: 208–215. doi:10.1016/j.scitotenv.2016.05.206

Lee, S. S., A. M. Paspalof, D. D. Snow, E. K. Richmond, E. J. Rosi-Marshall, and J. J. Kelly. 2016. Occurrence and potential biological effects of amphetamine on stream communities. Environ. Sci. Technol. 50: 9727–9735. doi:10.1021/acs.est.6b03717

Lemon, J. 2006. Plotrix: a package in the red light district of R. R-News 6: 8–12.

Leonhardt, S., and N. Blüthgen. 2012. The same, but different: pollen foraging in honeybee and bumblebee colonies. Apidologie 43: 449–464. doi:10.1007/s13592-011-0112-y

Lüdecke, D. 2018. ggeffects: Tidy Data Frames of Marginal Effects from Regression Models. J. Open Source Softw. 3: 772. doi:10.21105/joss.00772

Meador, J. P., A. Yeh, G. Young, and E. P. Gallagher. 2016. Contaminants of emerging concern in a large temperate estuary. Environ. Pollut. 213: 254–267. doi:10.1016/j.envpol.2016.01.088

Meyer, M. F., T. Ozersky, K. H. Woo, and others. 2022. Effects of spatially heterogeneous lakeside development on nearshore biotic communities in a large, deep, oligotrophic lake. Limnol. Oceanogr. 67: 2649–2664. doi:10.1002/lno.12228

Meyer, M. F., S. M. Powers, and S. E. Hampton. 2019. An evidence synthesis of pharmaceuticals and personal care products (PPCPs) in the environment: imbalances among compounds, sewage treatment techniques, and ecosystem types. Environ. Sci. Technol. 53: 12961–12973.

Pebesma, E. 2018. Simple Features for R: Standardized Support for Spatial Vector Data. R J. 10: 439–446. doi:10.32614/RJ-2018-009

Potts, S. G., J. C. Biesmeijer, C. Kremen, P. Neumann, O. Schweiger, and W. E. Kunin. 2010. Global pollinator declines: trends, impacts and drivers. Trends Ecol. Evol. 25: 345–353. doi:10.1016/j.tree.2010.01.007

R Core Team. 2022. R: A Language and Environment for Statistical Computing, R Foundation for Statistical Computing.

del Rey, Z. R., E. F. Granek, and B. A. Buckley. 2011. Expression of HSP70 in *Mytilus californianus* following exposure to caffeine. Ecotoxicology 20: 855–861. doi:10.1007/s10646-011-0649-6

Rheubert, J. L., M. F. Meyer, R. M. Strobel, M. A. Pasternak, and R. A. Charvat. 2020. Predicting antibacterial activity from snake venom proteomes. PloS One 15: e0226807.

Richmond, E. K., M. R. Grace, J. J. Kelly, A. J. Reisinger, E. J. Rosi, and D. M. Walters. 2017. Pharmaceuticals and personal care products (PPCPs) are ecological disrupting compounds (EcoDC). Elem Sci Anth 5: 66. doi:10.1525/elementa.252

Richmond, E. K., E. J. Rosi, A. J. Reisinger, B. R. Hanrahan, R. M. Thompson, and M. R. Grace. 2019. Influences of the antidepressant fluoxetine on stream ecosystem function and aquatic insect emergence at environmentally realistic concentrations. J. Freshw. Ecol. 34: 513–531. doi:10.1080/02705060.2019.1629546

Richmond, E. K., E. J. Rosi, D. M. Walters, J. Fick, S. K. Hamilton, T. Brodin, A. Sundelin, and M. R. Grace. 2018. A diverse suite of pharmaceuticals contaminates stream and riparian food webs. Nat. Commun. 9: 4491. doi:10.1038/s41467-018-06822-w

Shahriar, A., J. Tan, P. Sharma, D. Hanigan, P. Verburg, K. Pagilla, and Y. Yang. 2021. Modeling the fate and human health impacts of pharmaceuticals and personal care products in reclaimed wastewater irrigation for agriculture. Environ. Pollut. Barking Essex 1987 276: 116532. doi:10.1016/j.envpol.2021.116532

Slowikowski, K. 2019. ggrepel: Automatically Position Non-Overlapping Text Labels with “ggplot2.”

Thompson, H. M., and L. V. Hunt. 1999. Extrapolating from honeybees to bumblebees in pesticide risk assessment. Ecotoxicology 8: 147–166. doi:10.1023/A:1026444029579

Tong, Y., M. Wang, J. Peñuelas, and others. 2020. Improvement in municipal wastewater treatment alters lake nitrogen to phosphorus ratios in populated regions. Proc. Natl. Acad. Sci. 117: 11566–11572. doi:10.1073/pnas.1920759117

Walker, K. 2021. tigris: Load Census TIGER/Line Shapefiles.

Wei, T., and V. Simko. 2017. R package “corrplot”: Visualization of a Correlation Matrix.

Wickham, H., M. Averick, J. Bryan, and others. 2019. Welcome to the tidyverse. J. Open Source Softw. 4: 1686. doi:10.21105/joss.01686

Wickham, H., and J. Bryan. 2019. readxl: Read Excel Files.

Wilke, C. O. 2019. cowplot: Streamlined Plot Theme and Plot Annotations for “ggplot2.”

Wilkinson, J. L., A. B. A. Boxall, D. W. Kolpin, and others. 2022. Pharmaceutical pollution of the world’s rivers. Proc. Natl. Acad. Sci. 119: e2113947119. doi:10.1073/pnas.2113947119

Wright, G. A., D. D. Baker, M. J. Palmer, D. Stabler, J. A. Mustard, E. F. Power, A. M. Borland, and P. C. Stevenson. 2013. Caffeine in floral nectar enhances a pollinator’s memory of reward. Science 339: 1202–1204. doi:10.1126/science.1228806

