## Supplemental tables and figures for main manuscript for "Identifying drivers of sewage-associated pollutants in pollinators across urban landscapes"

| Table S1: Summary statistics of site data. Values represent cross-annual means, and values in parentheses represent cross-annual standard deviations. Site names are anonymized for confidentiality. | | | | | |
| --- | --- | --- | --- | --- | --- |
| **Site** | **Probability of PPCP Presence** | **Mean Development (%)** | **Mean Bee Abundance** | **Mean Bee Species Richness** | **Mean Plant Abundance** |
| site 1 | 0.33 | 58.81 (0.08) | 80.25 (18.54) | 4.70 (0.34) | 213.17 (84.42) |
| site 3 | 0.67 | 74.67 (0.42) | 77.16 (9.27) | 14.14 (5.36) | 447.80 (295.72) |
| site 4 | 0.14 | 82.00 (0.26) | 59.71 (7.53) | 9.58 (6.46) | 544.96 (270.03) |
| site 9 | 0.33 | 2.10 (0.00) | 51.29 (0.00) | 13.99 (0.00) | 1527.94 (0.00) |
| site 11 | 0.44 | 92.22 (0.01) | 56.87 (14.68) | 11.11 (4.76) | 878.49 (108.32) |
| site 12 | 0.67 | 30.47 (0.14) | 51.49 (39.19) | 12.97 (8.18) | 971.32 (15.93) |
| site 13 | 0.17 | 81.46 (0.57) | 76.33 (5.52) | 5.76 (0.79) | 165.98 (124.24) |
| site 15 | 0.00 | 49.07 (0.00) | 49.49 (0.00) | 4.71 (0.00) | 249.80 (0.00) |
| site 16 | 0.33 | 75.58 (0.00) | 66.83 (0.00) | 16.71 (0.00) | 662.24 (0.00) |
| site 18 | 0.55 | 91.91 (0.75) | 45.36 (5.20) | 10.79 (4.20) | 399.90 (303.63) |
| site 19 | 0.00 | 66.52 (0.00) | 62.93 (0.00) | 22.16 (0.00) | 519.40 (0.00) |
| site 21 | 0.00 | 15.08 (0.00) | 63.61 (0.00) | 16.31 (0.00) | 3422.59 (0.00) |
| site 24 | 0.33 | 52.17 (0.57) | 59.44 (15.09) | 5.90 (1.29) | 144.47 (26.30) |
| site 25 | 0.75 | 2.72 (0.14) | 123.49 (10.00) | 13.46 (5.43) | 4228.90 (6429.95) |
| site 26 | 0.33 | 10.93 (0.00) | 26.64 (0.00) | 5.04 (0.00) | 2017.75 (0.00) |
| site 34 | 0.17 | 60.82 (0.36) | 51.40 (7.86) | 5.46 (0.31) | 213.06 (90.59) |
| site 35 | 0.33 | 37.05 (0.76) | 113.40 (5.28) | 9.63 (0.96) | 592.37 (250.83) |

| Table S2: Summary statistics of taxon data. Values represent intra-specific, cross-site means, and values in parentheses represent intra-specific, cross-site standard deviations. | | | | | |
| --- | --- | --- | --- | --- | --- |
| **Species** | **Probability of PPCP Presence** | **Mean Development (%)** | **Mean Bee Abundance** | **Mean Bee Richness** | **Mean Plant Abundance** |
| *Agapostemon texanus* | 0.438 | 58.80  (29.27) | 67.43  (25.67) | 10.18  (5.58) | 655.02  (680.44) |
| *Apis mellifera* | 0.257 | 58.80  (30.36) | 68.50  (27.73) | 10.79  (5.78) | 1058.96  (2448.98) |
| *Bombus vosnesenskii* | 0.500 | 58.11  (30.53) | 68.93  (28.03) | 10.68  (5.83) | 1067.71  (2485.25) |

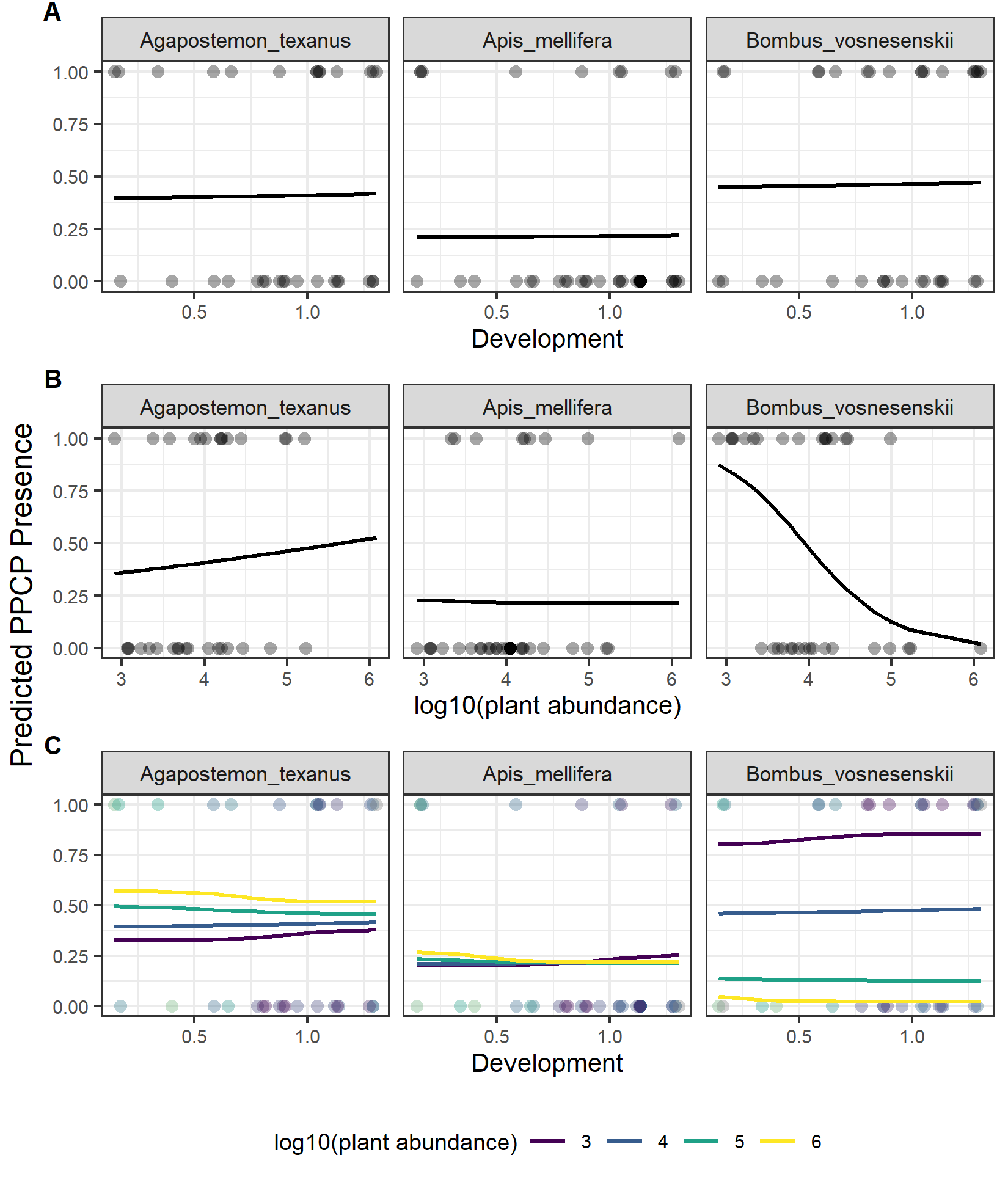

Figure S1: Probability of PPCP presence resulting from logistic regressions for human development (A), plant abundance (B), and the interaction of plant abundance and human development (C) by species. These two predictors and interaction terms were selected based on the most influential predictors from the model averaging routine.
