## Supplemental methods tables for main manuscript for "Identifying drivers of sewage-associated pollutants in pollinators across urban landscapes"

This document contains detailed information about PPCP extraction and HPLC/MS protocols. Tables referenced within this document are referenced as “Table SMXX”, where “SM” refers to “Supplemental Methods” and “XX” refers to the Table Number.

| Table SM1: List of PPCPs and associated information assessed in this study. | | | | | |
| --- | --- | --- | --- | --- | --- |
| Compound Name | Common Name | Use | CAS Number of standards used in this analysis | Selected-ion monitoring ion masse, in mass/charge | Time (minutes) |
| 11,7-Dimethylxanthine | Paraxanthine | Metabolite of caffeine (stimulant) | 611-59-6 | 124.1, 181.0 | 3.79 |
| Acetaminophen | Tylenol | Analgesic | 103-90-2 | 110.1, 152.0 | 2.68 |
| Caffeine | - | Stimulant | 58-02-8 | 138.0, 195.1 | 9.10 |
| Cotinine | - | Metabolite of nicotine | 486-56-6 | 80.1, 98.1, 177.1 | 2.60 |
| Diphenhydramine | Benadryl | Antipruritic | 58-73-1 | 167.1,256.1 | 19.45 |
| Sulfamexthazole | - | Antibiotic | 723-46-6 | 108.0, 254.0 | 16.24 |
| Warfarin | Coumadin | Anticoagulant; rodenticide | 81-81-2 | 163.0, 309.1 | 23.11 |

| Table SM2: Timetable of liquid reagent used throughout HPLC/MS quantification. | | |
| --- | --- | --- |
| Time (minutes) | Percentage of 10-mM formate buffer | Percentage of 100% acetonitrile |
| 0 | 94 | 6 |
| 5 | 94 | 6 |
| 9 | 86 | 14 |
| 10 | 76 | 24 |
| 15 | 59 | 41 |
| 16 | 49 | 51 |
| 26 | 30 | 70 |
| 27 | 0 | 100 |
| 39 | 0 | 100 |
| 45 | 94 | 6 |
| 65 | 94 | 6 |

| Table SM3: HPLC/MS settings used for sample processing in the HPLC/MS. | |
| --- | --- |
| Characteristic | Setting |
| Nitrogen dry gas temperature: | Nitrogen dry gas temperature: |
| 350 degrees Celsius | 350 degrees Celsius |
| Drying gas flow rate: | Drying gas flow rate: |
| 12.0 liters per minute | 12.0 liters per minute |
